# Gene knockout shows that PML (TRIM19) does not restrict the early stages of HIV-1 infection in human cell lines

**DOI:** 10.1101/109769

**Authors:** Nasser Masroori, Pearl Cherry, Natacha Merindol, Jia-xin Li, Caroline Dufour, Lina Poulain, Mélodie B. Plourde, Lionel Berthoux

## Abstract

The PML (promyelocytic leukemia) protein is a member of the TRIM family, a large group of proteins that show high diversity in functions but possess a common tripartite motif giving the family its name. We and others recently reported that both murine PML (mPML) and human PML (hPML) strongly restrict the early stages of infection by HIV-1 and other lentiviruses when expressed in mouse embryonic fibroblasts (MEFs). This restriction activity was found to contribute to the type I interferon (IFN-I)-mediated inhibition of HIV-1 in MEFs. Additionally, PML caused transcriptional repression of the HIV-1 promoter in MEFs. By contrast, the modulation of the early stages of HIV-1 infection of human cells by PML has been investigated by RNAi with unclear results. In order to conclusively determine whether PML restricts HIV-1 or not in human cells, we used CRISPR-Cas9 to knock out its gene in epithelial, lymphoid and monocytic human cell lines. Infection challenges showed that PML knockout had no effect on the permissiveness of these cells to HIV-1 infection. IFN-I treatments inhibited HIV-1 equally whether PML was expressed or not. Over-expression of individual hPML isoforms, or of mPML, in a human T cell line did not restrict HIV-1. The presence of PML was not required for the restriction of nonhuman retroviruses by TRIM5α was inhibited by arsenic trioxide through a PML-independent mechanism. We conclude that PML is not a restriction factor for HIV-1 in human cell lines representing diverse lineages.

**Importance:** PML is involved in innate immune mechanisms against both DNA and RNA viruses. Although the mechanism by which PML inhibits highly divergent viruses is unclear, it was recently found that it can increase the transcription of interferon-stimulated genes (ISGs). However, whether human PML inhibits HIV-1 has been debated. Here we provide unambiguous, knockout-based evidence that PML does not restrict the early post-entry stages of HIV-1 infection in a variety of human cell types and does not participate in the inhibition of HIV-1 by IFN-I. Although this study does not exclude the possibility of other mechanisms by which PML may interfere with HIV-1, we nonetheless demonstrate that PML does not generally act as an HIV-1 restriction factor in human cells and that its presence is not required for IFN-I to stimulate the expression of anti-HIV-1 genes. These results contribute to uncovering the landscape of HIV-1 inhibition by ISGs in human cells.

## Introduction

PML/TRIM19 belongs to the tripartite motif (TRIM) protein superfamily that shares a conserved tripartite architecture: a RING domain, one or two B-boxes, and a coiled-coil domain (1). Due to the alternative splicing of the C-terminal domain, seven PML isoforms are present in human cells. Isoforms I to VI are primarily located in the nucleus, while PML VII is mostly cytoplasmic (2). PML is the major component of a nuclear substructure named PML nuclear body (PML NB). PML NBs are dynamic and their size, number, and composition change in response to cellular stresses or during the cell cycle. In addition to PML, these NBs recruit many other proteins in a transient fashion (3–6). TRIM5α, a cytoplasmic factor that restricts retroviruses in a species-specific, virus-specific manner (7), is actively shuttling between the cytoplasm and the nucleus and localizes to the PML NBs when present in the nucleus (8). PML is involved in many cellular activities including transcriptional regulation and tumor suppression (5, 9, 10).

IFNs are a multigene family of inducible cytokines released by host cells in response to pathogens, including viruses (11–13). IFN-I binding to its receptor leads to the transcriptional stimulation of a set of genes encoding antiviral proteins which inhibit the replication of a wide range of viruses (12, 14). The transcription of PML and of many NB-associated proteins (e.g. Daxx and Sp100) is up-regulated by IFN-I (15, 16). Conversely, it was recently proposed that PML is involved in the IFN-I-induced expression of ISGs by directly binding their promoter (17).

The involvement of PML in antiviral defense mechanisms against several DNA and RNA viruses has been extensively studied. PML was shown to restrict a complex retrovirus, the human foamy virus, by inhibiting viral gene expression (18). PML deficient cells are also more prone to infection with rabies virus (19). Moreover, PML was shown to interfere with the replication of poliovirus (20), encephalomyocarditis virus (EMCV) (21), herpes simplex virus type-1 (HSV-1), adeno-associated virus (AAV) (22), influenza virus, and vesicular stomatitis virus (VSV) (23). As a direct consequence, some viruses such as HSV-1 and the human cytomegalovirus have evolved mechanisms to counteract PML, either by disrupting PML NBs and/or by inducing PML degradation (24–26).

The role of PML in HIV-1 infection of human cells is controversial. As_2_O_3_, a drug that induces PML oligomerization and degradation (27), was shown to increase the susceptibility of human cells to N-tropic murine leukemia virus (N-MLV) and HIV-1 (28). A recent study proposed that PML was an indirect inhibitor of HIV-1 early post-entry infection stages through its association with Daxx, a constitutive partner protein in PML NBs (29). However, another group found that the depletion of PML (but not that of Daxx) enhanced HIV-1 infection in human primary fibroblasts, while having no effect in T cell lines such as Jurkat (30). PML was also found to regulate HIV-1 latency. Specifically, PML degradation or NBs disruption resulted in the activation of HIV-1 provirus transcription in a lymphoid model of HIV-1 latency (31), although these results have not been independently confirmed. There is consensus, however, on the existence of a PML-dependent restriction of HIV-1 in MEFs. In these cells, PML inhibits the early post-entry stages of infection (32–34) and also promotes the transcriptional silencing of the integrated provirus (34). Human PML (hPML) was able to reconstitute both restriction activities in MEFs knocked out for the endogenous murine PML (mPML), in an isoform-specific fashion (34). In addition, the inhibition of lentiviruses by IFN-I in MEFs involves PML (34). In this study, we investigate the role of PML in the restriction of HIV-1 and other retroviruses in several human cell lines, including T cells and myeloid cells, by gene knockout. We also examine the role of PML in the IFN-induced restriction of lentiviruses in human cells. We show that PML is dispensable for the restriction of lentiviruses in human cells, is not involved in the IFN-I-mediated inhibition of infection, and is not relevant to the inhibition of TRIM5α by As_2_O_3_.

## Results

### CRISPR-Cas9-mediated knockout of PML in human cells

In order to stably and irreversibly deplete PML in human cells, we designed two guide RNAs (gRNAs), hPML1 and hPML2, to target the Cas9 nuclease towards exon 2 of *PML* (Fig. 1). Exon 2 is present in all hPML isoforms, and the algorithm used to design the gRNAs minimizes the risk of nonspecific targeting. The plasmid used in this study, pLentiCRISPRv2 (pLCv2), can mediate knockouts through transfection and also through lentiviral transduction. The control plasmid, pLCv2-CAG, targets the CMV-IE/chicken actin/rabbit beta globin hybrid promoter, a nonhuman sequence (35). We used the Surveyor assay (36) to reveal the presence of insertions/deletions (indels) in the PML gene of HEK293T cells transiently transfected with pLCv2-hPML1 or pLCv2-hPML2. We could observe the presence of PML DNA digestion products of the expected size in cells transfected with each of the PML gRNAs but not in cells transfected with the control gRNA (Fig. 1A), indicating that both PML gRNAs generated double-strand breaks that were repaired by non-homologous end joining (NHEJ). To quantify the extent of DNA damage following stable lentiviral transduction of the CRISPR components, we transduced human monocytic THP-1 cells with the LCv2-hPML1 vector and, as a control, the irrelevant LCv2-CAG vector. Cells were treated with puromycin to eliminate non-transduced cells, and amplicons of the targeted PML region were then obtained and Sanger sequenced. A reference contig alignment of the sequencing plots revealed that a −1 deletion was the most prevalent mutation found in LCv2-hPML1-transduced cells, but other types of indels were present, as evidenced by the presence of additional peaks at each position (Fig. 1B). We further analyzed the sequencing data using the Tracking of Indels by Decomposition (TIDE) method available online (see Methods) (Fig. 1C). Computations using this assay showed that at least 96.3% of PML alleles contained an indel at the expected position in cells transduced with the hPML1 gRNA.

**FIG 1.**
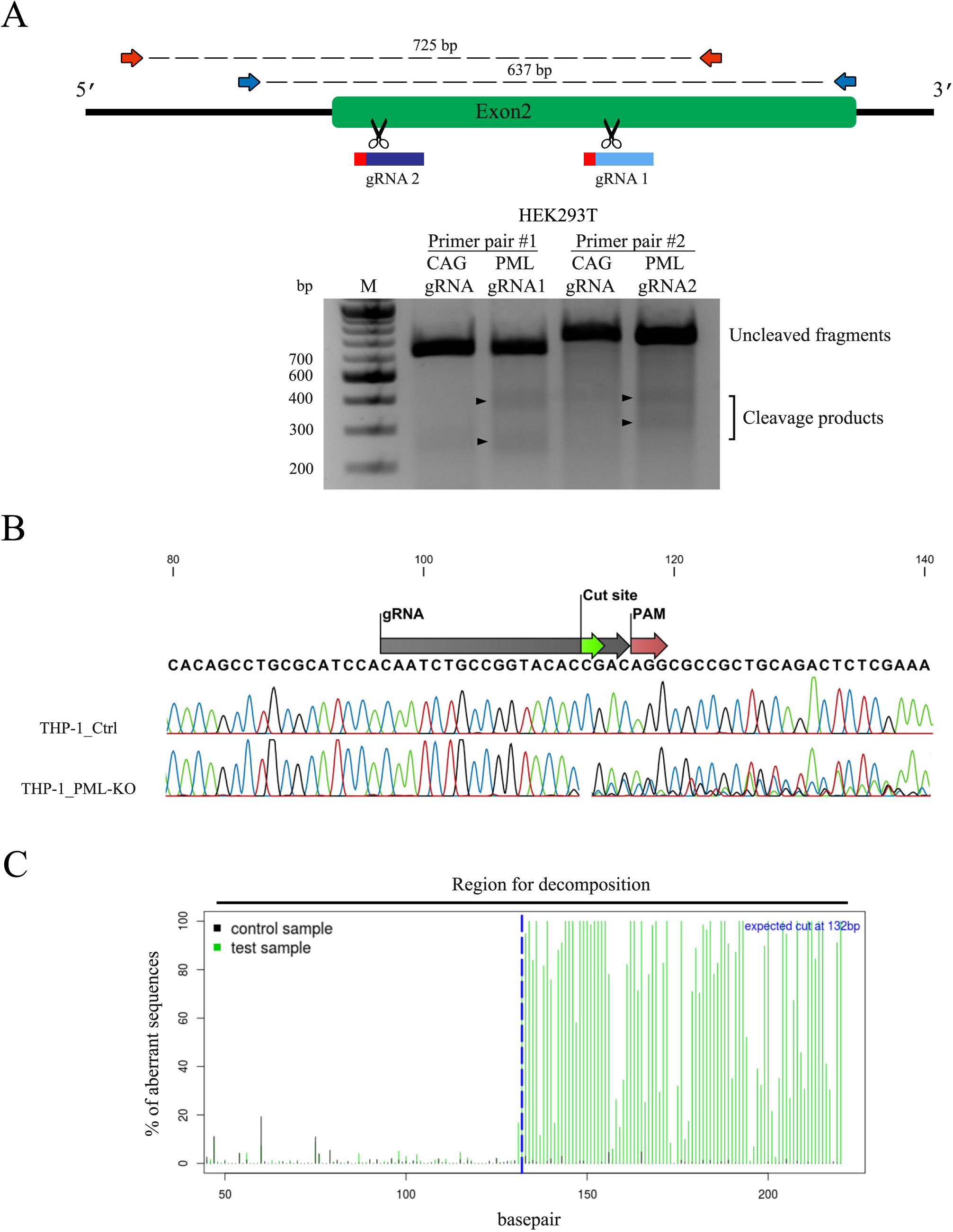
CRISPR/Cas9-mediated genome editing of PML in human cell lines. (A) The Cas9 nuclease was targeted to exon 2 of the PML gene (green) by two selected gRNAs whose binding sites are shown in blue (PAM motifs are in red). Arrows indicate the positions of the binding sites for the ODNs used in the PCR-Surveyor assay (blue arrows for gRNA1-, red arrows for gRNA2-guided cut sites). The Surveyor assay is shown in the lower panel. Briefly, PCR products amplified from cells transfected with pLCv2-hPML1, -hPML2, or pLCv2-CAG (Ctrl), were subjected to denaturation, re-annealing and digestion with the Surveyor enzyme. Arrowheads indicate cleavage products of the expected size. (B) Sanger sequencing analysis of *PML* in cells transduced with LCv2-hPML1. THP-1 cells were transduced with lentiviral vectors produced using pLCv2-hPML1 or the control pLCv2-CAG. Following puromycin selection, the targeted *PML* locus was PCR-amplified and the PCR product was Sanger sequenced. The figure shows an alignment of the obtained sequence plots. (C) Decomposition of sequencing plots by TIDE assay. The graph shows the % of aberrant peaks upstream and downstream of the cut site in the sequencing reactions shown in panel B. The % of indel-containing alleles was computed by TIDE.

### Knocking out PML in human monocytic cells has little to no effect on the permissiveness to HIV-1 in the presence or absence of IFN-I

THP-1 cells were stably transduced with lentiviral vectors produced using pLCv2-hPML1 and pLCv2-hPML2. Following puromycin selection, we performed a Western blotting (WB) analysis of PML levels in bulk populations (Fig. 2A). The levels of hPML were not sufficiently high to be detected in unstimulated cells (not shown), and therefore, the analysis was done using cells treated with IFN-β. In control cells we found several bands corresponding to hPML isoforms, as previously reported (2). In the cells transduced with the hPML gRNAs, PML was undetectable, showing that knockout was efficient with both gRNAs and affected all detectable isoforms. This result is consistent with the NHEJ-mediated mutagenesis observed in transfected HEK293T cells using both gRNAs shown in Fig. 1. As both gRNAs showed similar efficiency, all the subsequent experiments in this study were only performed with one gRNA, hPML1. We next infected PML knockout (hPML1 gRNA transduced) and control cells (CAG transduced) with a single dose of HIV-1_NL-GFP_ (37), a VSV-G-pseudotyped, Δ-Envelope HIV-1 vector expressing GFP in place of Nef (Fig. 2B). The percentage of GFP-positive cells following HIV-1_NL-GFP_ challenge is directly proportional to the cells’ permissiveness toward infection by this virus. This system is thus well-suited to analyze restriction activities taking place during post-entry steps and until integration. These infections were performed in the presence or absence of IFN-β, owing to the reported role of PML in stimulating the transcription of ISGs (38). In the absence of IFN-β, we found that the PML-KO cells were slightly more permissive to infection by HIV-1_NL-GFP_ compared with the control cells (less than 2-fold). The addition of IFN-β very strongly inhibited (>20-fold) the infection of THP-1 cells (Fig. 2B), and the low numbers of infected cells prevented a fine analysis of the role of PML in this inhibition. However, the absence of PML clearly did not prevent IFN-β from inhibiting HIV-1_NL-GFP_, showing that PML was dispensable for this activity.

**FIG 2.**
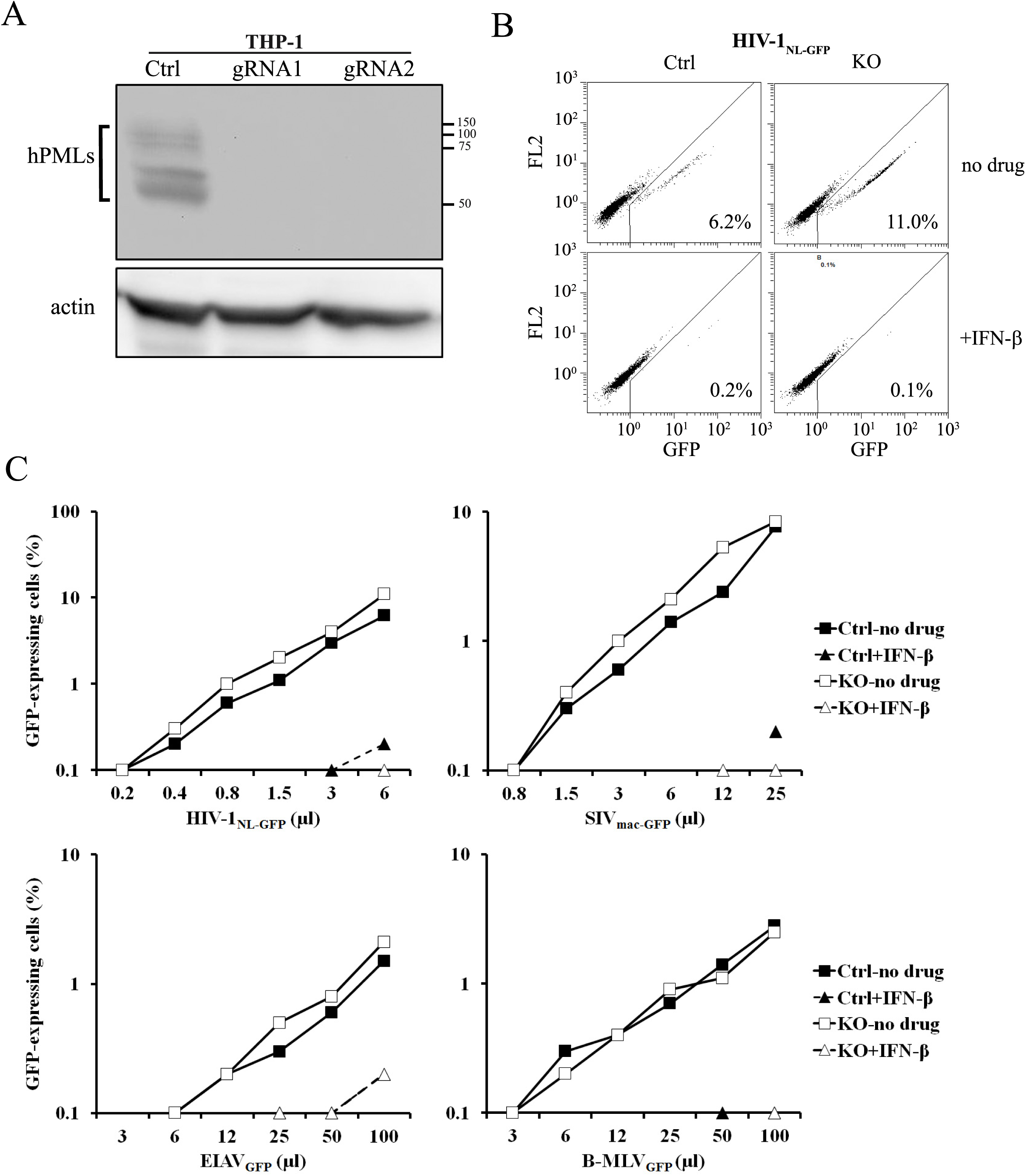
PML knockout has negligible effects on intrinsic or IFN-I-induced restriction of retroviruses in THP-1 cells. (A) WB analysis of THP-1 cells transduced with pLCv2-based vectors expressing Cas9 and a gRNA targeting either hPML or CAG. Stably transduced, puromycin-resistant cells were treated with IFN-β (10 ng/ml). Cellular lysates were prepared 16 h later and analyzed by WB using antibodies against hPML and actin as a loading control. (B) FACS plots from PML-knockout (KO) and control (Ctrl, CAG gRNA-transduced) THP-1 cells infected with HIV-1_NL-GFP_. Control or PML-KO THP-1 cells were treated with IFN-β or left untreated and then exposed to HIV-1_NL-GFP_ (10 µl). 2 d later, cells were analyzed by FACS and the percentage of infected (GFP-positive) cells observed is indicated on each plot. (C) Virus dose-dependent analysis of the role of hPML in the intrinsinc and IFN-I-induced restriction of retroviruses. Control and PML-KO THP-1 cells were treated with IFN-β (10 ng/ml) for 16 h, followed by infection with increasing doses of retroviral vectors. The percentage of infected cells was assessed 2 d later by FACS.

In order to obtain a more complete picture of the importance of PML in the permissiveness to retroviruses in this immune cell line, we performed additional infections with this HIV-1 vector as well as with GFP-expressing vectors derived from the macaque strain of the simian immunodeficiency virus (SIV_mac-GFP_), the equine infectious anemia virus (EIAV_GFP_) and the B-tropic murine leukemia virus (B-MLV_GFP_). EIAV is restricted by TRIM5α in human cells (39), making it possible to analyze whether PML modulates the restriction of retroviruses by this well-characterized restriction factor. Infectivity of the three lentiviral vectors (HIV-1, SIVmac, EIAV) was slightly higher in the absence of PML at most virus doses used, whereas infectivity of the B-MLV vector was unaffected by PML knockout (Fig. 2C). These results suggest that PML has a small, barely detectable inhibitory effect on the infection of THP-1 cells by lentiviruses and does not modulate TRIM5α activity. Treatment with IFNβ-strongly decreased THP-1 permissiveness to all four vectors, preventing us from measuring the -fold decrease in infectivity with accuracy (Fig. 2C). However, it was clear that IFN-β efficiently inhibited infection in the presence or absence of PML, indicating that PML is not crucial for the IFN-Imediated anti-retroviral response.

### Knocking out PML in human epithelial cells has little to no effect on the permissiveness to retroviral infections in the presence or absence of IFN-I

We then transduced epithelial carcinoma HeLa cells with the CAG or PML gRNAs. PML was efficiently knocked out, as seen by WB (Fig. 3A). We also performed immunofluorescence microscopy to analyze the effect of PML knockout on PML and SUMO nuclear bodies. A large part (but not all) of SUMO-1 localizes to PML bodies in normal conditions (40). As expected, signal corresponding to PML nuclear bodies almost completely disappeared from the cells transduced with the PML gRNA (Fig. 3B). In addition, SUMO-1 punctate nuclear staining was strongly diminished but not abolished (Fig. 3B). We then challenged the stably transduced cells with GFP-expressing viral vectors like we had done in THP-1 cells. We found that susceptibility to HIV-1, SIVmac, EIAV and B-MLV vectors was identical whether PML was present or not (Fig. 3C-D). IFN-β inhibited all four viral vectors, although the magnitude of this effect (∼2- to 3-fold) was much smaller than in THP-1 cells. IFN-β treatments had identical effects in PML-expressing and PML-knockout cells, again showing that PML does not modulate this inhibitory pathway in human cells.

**FIG 3.**
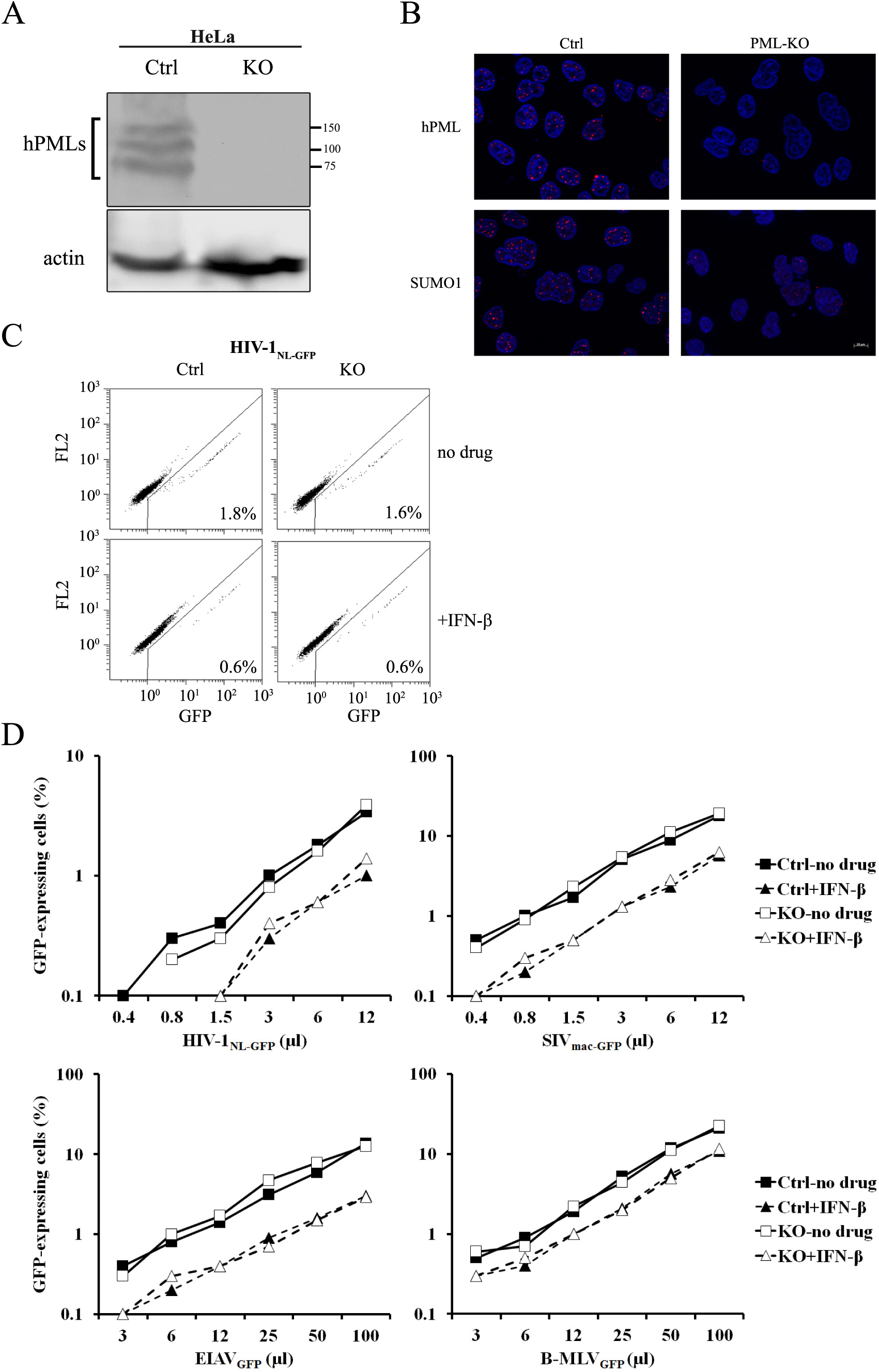
PML knockout has no effect on intrinsic or IFN-I-induced restriction of retroviruses in HeLa cells. (A) HeLa cells lentivirally transduced with pLCv2 vectors expressing either the hPML gRNA1 or (as a control) the CAG-targeting gRNA were treated with IFN-β (10 ng/ml). Cellular lysates were prepared 16 h later and analyzed by WB using an anti-hPML antibody. Actin was analyzed as a loading control. (B) IF microscopy analysis of PML bodies in HeLa cells transduced with LCv2-PML1 (PML-KO) or transduced with LCv2-CAG as a control (Ctrl). Puromycin-selected cells were stained for PML (top) or SUMO-1 (bottom). Nuclei were stained with Hoechst33342. (C) FACS plots from transduced HeLa cells infected with HIV-1_NL-GFP_. Control and PML-KO HeLa cells treated or not with IFN- were infected with HIV-1_NL-GFP_ (6 µl). The percentage of infected cells determined at 2 d post-infection is indicated for each plot. (D) Virus dose-dependent analysis of the role of hPML in IFN-I-induced restriction of retroviral infection. Control and PML-KO HeLa cells were treated with IFN- β, followed 16 h later by infection with increasing doses of the indicated retroviral vectors. The percentage of infected cells was assessed 2 d later by FACS.

Rhabdomyosarcoma-derived epithelial TE671 cells were similarly knocked out for PML by lentiviral transduction, and knockout was efficient (Fig. 4A). Similar to what we found in HeLa cells, infectivity of the four vectors tested was identical whether PML was present or not (Fig. 4B). IFN-β decreased the permissiveness of TE671 cells to all four vectors, although we noticed that IFN-β had a relatively smaller effect on HIV-1_NL-GFP_ compared with the three other vectors in TE671 (Fig. 4B). The IFN-β-induced inhibition of the four retroviral vectors in TE671 cells was identical whether PML was present or not (Fig. 4B).

**FIG 4.**
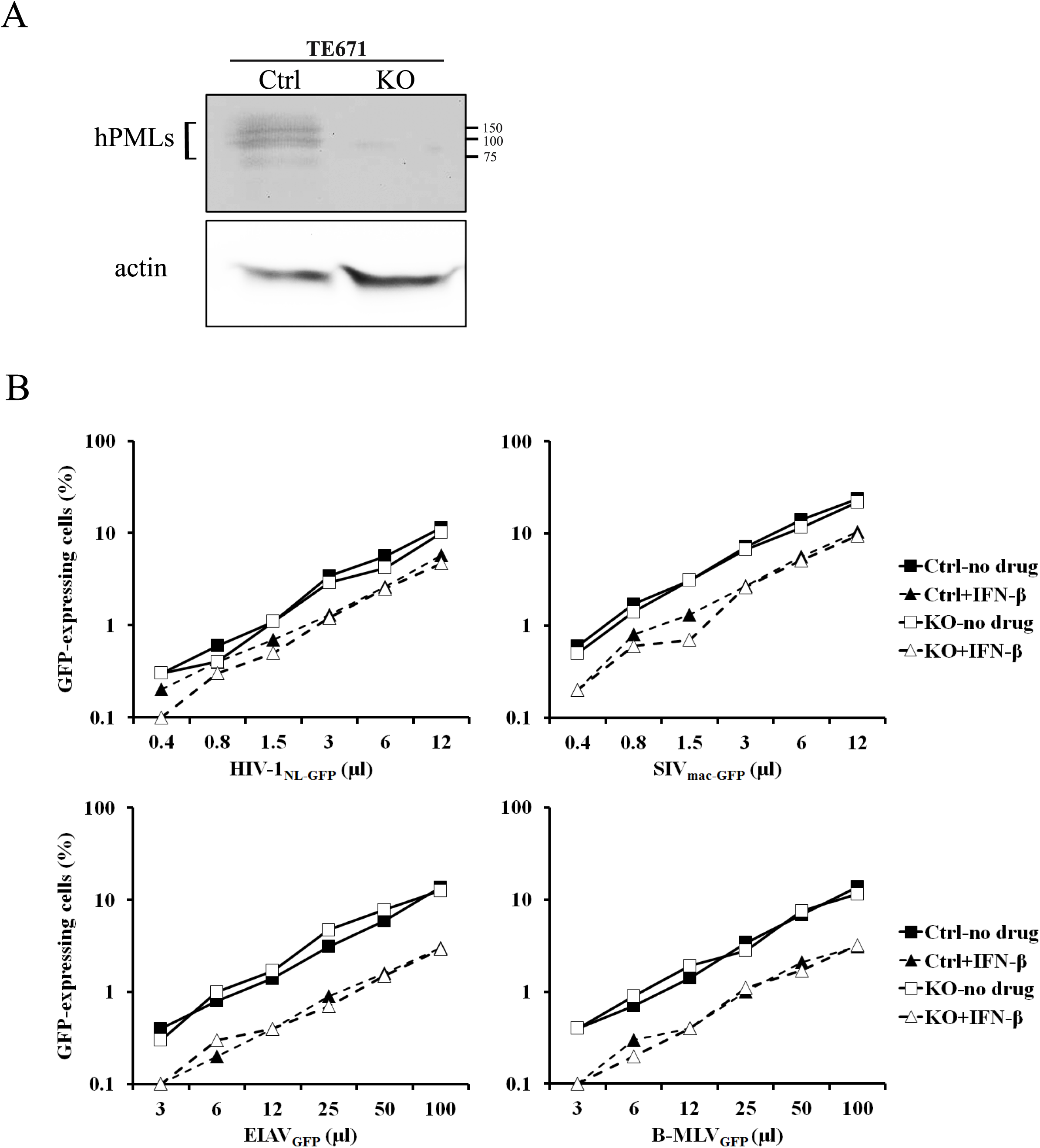
PML knockout has no effect on intrinsic or IFN-I-induced restriction of HIV-1 in TE671 cells. (A) WB analysis. TE671 cells were stably transduced with pLCv2-based vectors expressing Cas9 and either the hPML-targeting gRNA1 or the CAG-targeting gRNA as a control. The cells were treated with IFN-β (10 ng/ml) or left untreated as a control. Cellular lysates were prepared 16 h later and analyzed by WB using an anti-hPML antibody along with actin as a loading control. (B) Infection assay. Control (CAG gRNA-transduced) and PML-KO TE671 cells were treated with IFN- β or left untreated. 16 h later, the cells were infected with increasing doses of the indicated retroviral vectors. The percentage of infected cells was assessed 2 d later by FACS.

### A knock-in approach to suppress PML in human cells

In order to achieve efficient knockout by transient transfection without the need to isolate cellular clones by limiting dilution, we constructed a plasmid to serve as donor DNA in homology-directed repair (HDR). This plasmid contains two ∼800bp-long PML homology arms surrounding a neomycin resistance cassette (Fig. 5A). It is expected that its co-transfection in cells along with Cas9 and the hPML gRNA1 would lead to the knock-in of Neo^R^ in *PML* through HDR in a fraction of the cells. Selection in neomycin then eliminates cells in which the knock-in did not occur. Even if not all alleles of a given gene are successfully modified by knock-in, recent reports indicate that the remaining ones are usually knocked out through NHEJ-dependent mechanisms (41). We designed PCR primers for the specific amplification of the knock-in product and another pair to amplify the wild-type (WT) or the NHEJ-repair knockout alleles (Fig. 5A). To validate this system, we co-transfected TE671 cells with pLCv2.PML1 and the HDR donor plasmid, and randomly isolated a number of neomycin-resistant cell clones of which a representative analysis is shown in Fig. 5B. The knock-in product was detected as expected in all 7 clones while being absent in the parental cells. On the other hand, the band corresponding to WT or NHEJ-repaired alleles was less intense in these clones relative to the parental cells, but was always present, suggesting that HDR-mediated knockout did not affect all the *PML* alleles.

**FIG 5.**
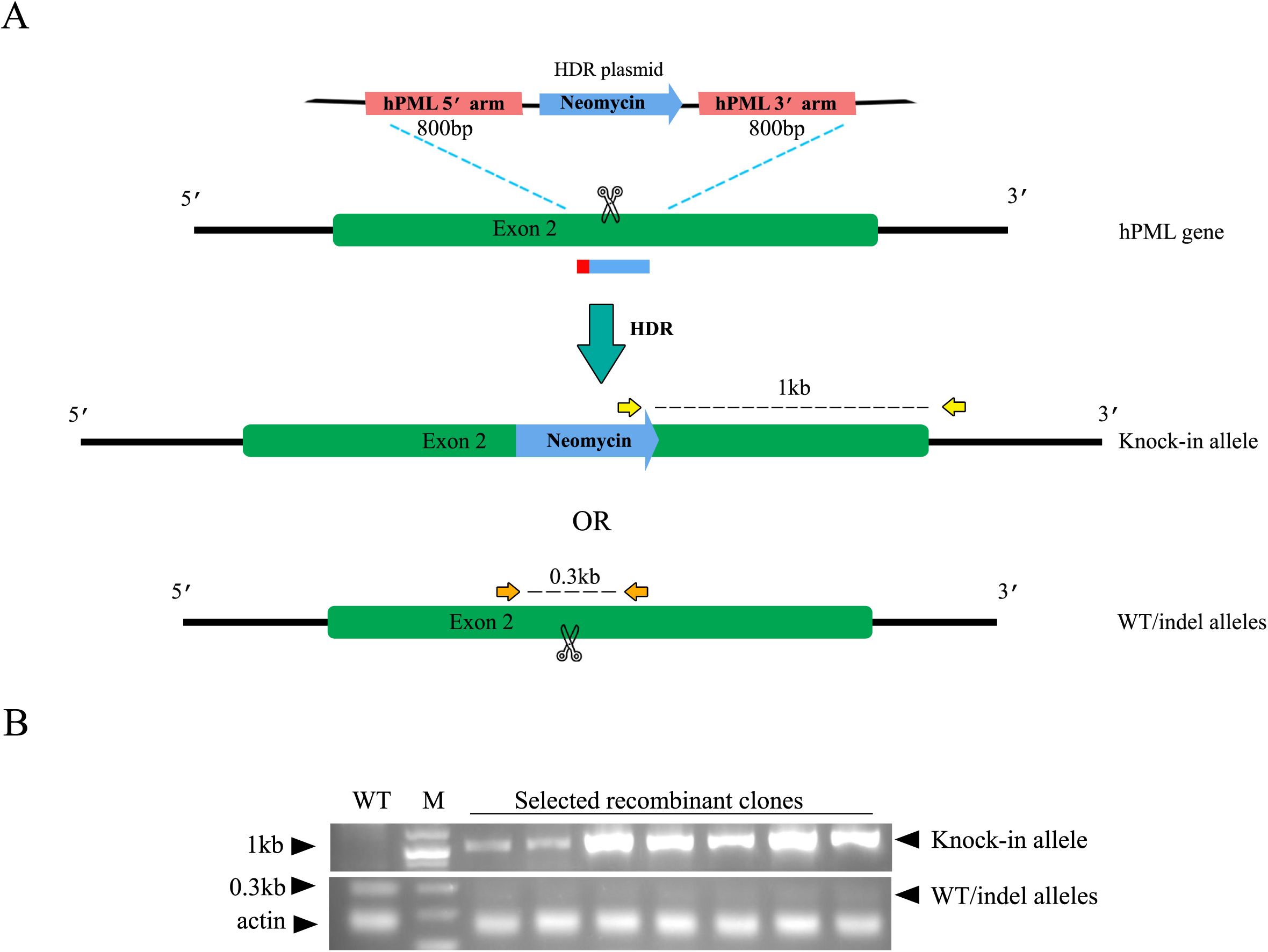
HDR-mediated knockout of PML. (A) Schematic of the HDR plasmid and targeting strategy for the knock-in of the Neomycin resistance gene at the PML locus. Two 800 bp-long PML homology arms encompass the Neo^R^ expression cassette on plasmid pNMs-Neo.HDR-hPML. The arms are complementary to the PML regions on either side of the gRNA1-mediated Cas9 cleavage site. Co-transfection of pLCv2-hPML1 and pNMs-Neo.HDR-hPML may yield a knock-in allele as indicated if DNA is repaired by HDR. If DNA is repaired by NHEJ, WT or indel-containing alleles may be generated. Yellow and orange arrows indicate the binding sites for the primers used to detect knock-in and WT/indel alleles by PCR (1 Kbp and 0.3 Kbp products, respectively). (B) PCR analysis of Neomycin-resistant TE671 clones. TE671 cells were co-transfected with pLCv2-hPML1 and pNMs-Neo.HDR-hPML, then grown in presence of neomycin. Individual Neo^R^ clones were analyzed using the two primer pairs described in A.

### PML is important for the efficient inhibition of SIVmac but not HIV-1 by IFN-I in lymphoid cells

We knocked out PML in Jurkat cells using the transfection approach that results in the insertion of *Neo^R^* in *PML*, as described above. We performed WB analyses to assess knockout efficiency. (Fig. 6A). Treatment with the IFN-I species IFN-α, IFN-β and IFN-ω stimulated PML expression in Jurkat cells. PML was efficiently knocked out (Fig. 6A), validating the HDR-based approach. In the absence of IFN-β, PML had little effect on the infectivity of all four vectors (□2-fold) (Fig. 6B). The effect of IFN-β treatment differed according to the retroviral vector used (Fig. 6B). IFN-β treatment decreased HIV-1_NL-GFP_ infectivity by ∼3.5-fold in both control and PML-KO cells. IFN-β similarly decreased the infectivity of SIV_mac-GFP_ by about 4-fold, but only in the control cells. In the PML-KO cells, the inhibitory effect of IFN-β on SIV_mac-GFP_ infectivity was smaller (□2-fold). Interestingly, we found the opposite situation upon challenge with the EIAV_GFP_ vector: IFN-βtreatment had no effect on EIAV_GFP_ infectivity in the WT Jurkat cells, whereas it significantly inhibited this vector in PML-KO cells, especially at low vector doses. Finally, IFN-β decreased the infectivity of B-MLV_GFP_ in both WT cells and PML-KO cells, with no apparent specificity. Thus, Jurkat cells provided a more complex situation with respect to the importance of PML in the antiviral effects. In order to further study the contrasting phenotypes of the HIV-1 and SIVmac vectors in these cells, we also analyzed the effects of IFN-α and IFN-ω. (Fig. 6C). We found that in control cells, all three IFN-I species decreased infectivity of both the HIV-1 and the SIVmac vectors, by 2- to 4-fold; IFN-β appearing to be the most consistently inhibitory IFN-I in these cells, similar to what we had observed in other cell lines (not shown) and to what was reported in the literature (42). In PML-KO cells, HIV-1_NL-GFP_ was inhibited by all three IFN-I species, similar to the control cells. In contrast, IFN-I inhibition of SIV_mac-GFP_ was much less efficient in PML-KO cells (Fig. 6C, bottom right panel). Thus, PML is important for IFN-I to inhibit the early infection stages of SIVmac, but not HIV-1, in Jurkat cells.

**FIG 6.**
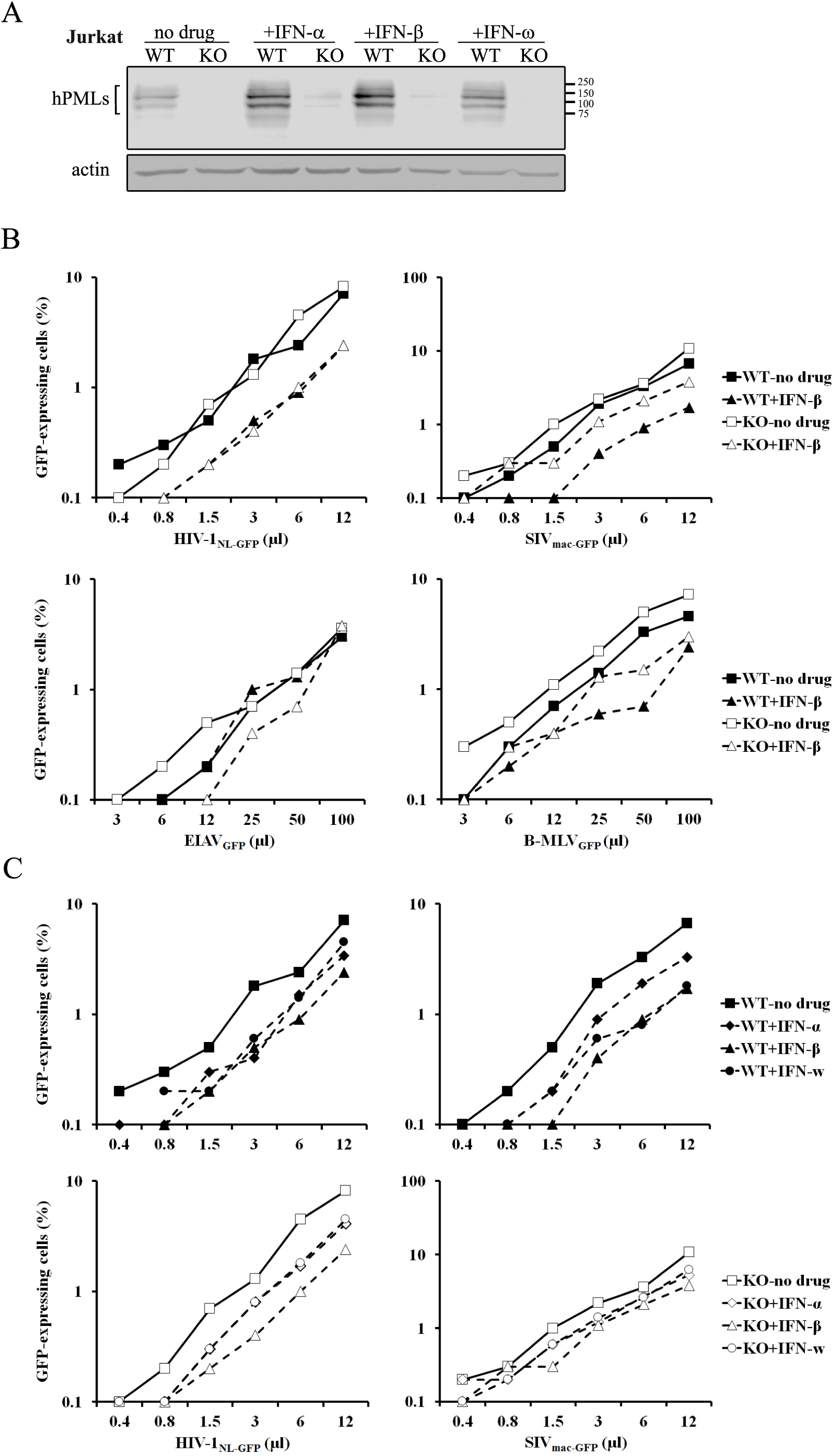
PML knockout has virus-specific effects on the restriction of retroviruses in Jurkat cells. (A) Jurkat cells were co-transfected with pLCv2.hPML1 and pNMs-Neo.HDR-hPML. Neomycin-resistant cells (KO) and parental untransfected cells (WT) were treated with IFN-α IFN- β or IFN-ω(10 ng/ml). Cellular lysates were prepared 16 h later and analyzed by WB using an anti-hPML antibody. Actin was analyzed as a loading control. (B) Virus dose-dependent analysis of the role of hPML in the intrinsic and IFN-I-induced restriction of retroviruses. PML-KO and control Jurkat cells were treated with IFN-β for 16 h, followed by infection with increasing doses of the indicated retroviral vectors. The percentage of infected cells was assessed 2 d later by FACS. (C) PML-KO and control cells were challenged with increasing doses of HIV-1_NL-GFP_ following treatment with IFN-α, −β or –ω for 16 h. The percentage of infected cells was assessed 2 d later by FACS.

### Over-expression of murine or human PML in Jurkat cells does not affect the infectivity of an HIV-1 vector

Unlike the PML-KO THP-1, HeLa and TE671 cells, the PML-KO Jurkat cells generated do not continuously express Cas9 or a PML-targeting gRNA. Thus, these cells provided an appropriate model to test whether the over-expression of specific hPML isoforms in a PML-KO background could inhibit HIV-1 or other retroviruses. In other words, this experiment was designed to reveal a possible cryptic restriction activity associated with specific PML isoforms that would normally not be apparent due to the presence of other isoforms. We retrovirally transduced the isoforms I to VI of hPML into PML-KO Jurkat cells, separately.

Because HIV-1 is inhibited by mPML in MEFs (32–34), we also transduced mPML. A WB analysis showed that all six isoforms of hPML were expressed, as was mPML isoform 2 (Fig. 7A). We then challenged the various cell cultures with the HIV-1, SIVmac, EIAV and B-MLV vectors (Fig. 7B). We found that none of the PML isoforms had an effect on GFP transduction by HIV-1_NL-GFP_. Interestingly, several hPML isoforms and mPML slightly increased permissiveness to SIV_mac-GFP_, by ∼2-fold. Permissiveness to EIAV_GFP_ was overall not modulated by over-expression of hPML or mPML, although a slight increase in infectivity was observed in presence of some hPML isoforms at the highest virus doses tested. Finally, the presence of hPML-VI slightly inhibited infection by B-MLV_GFP_ at least at some virus doses used (Fig. 7B). Thus, although individual PML isoforms modestly modulated the permissiveness to infection by the SIVmac, EIAV and B-MLV vectors in a virus-specific fashion, none of them affected permissiveness to infection by the HIV-1 vector.

**FIG 7.**
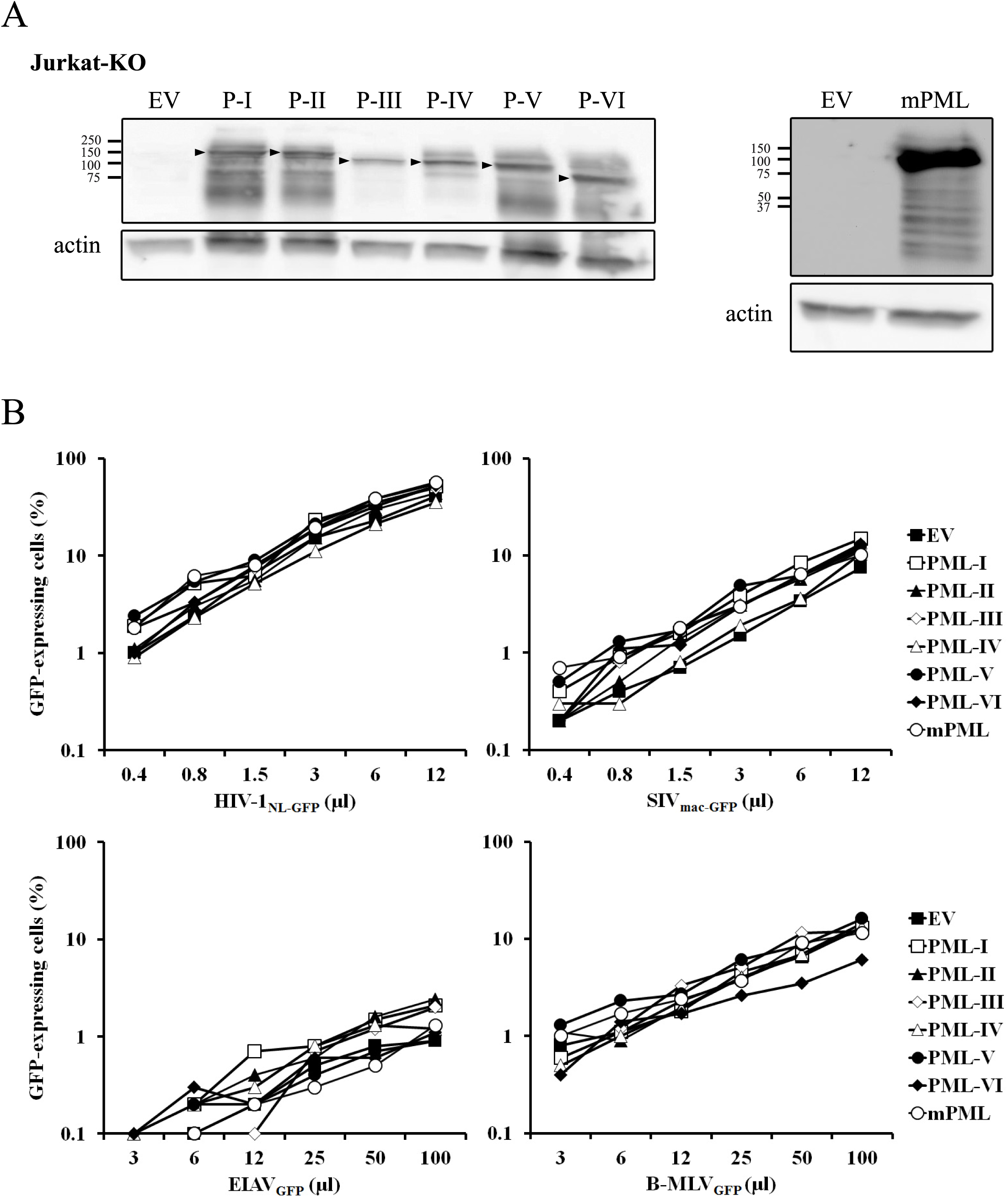
Transduction of mPML or hPML isoforms in PML-KO Jurkat cells has virus-specific effects on the permissiveness to retroviral vectors. (A) WB analysis of mPML and hPML expression. PML-KO Jurkat cells were stably transduced with mPML or with FLAG-tagged hPML-I to -VI separately. The empty vector (EV) was transduced as a control. Lysates prepared from the different cell populations were analyzed by WB with anti-FLAG (left panel) or anti-mPML (right panel) antibodies. Actin was probed as loading control. The arrowheads indicate the expected size for each hPML isoform. (B) Susceptibility to transduction by retroviral vectors. The cells were infected with multiple doses of the indicated retroviral vectors and the percentage of GFP-expressing cells was determined 2 d later by FACS.

### Restriction of N-MLV by TRIM5α and inhibition of TRIM5α by arsenic trioxide are independent of PML

Intriguingly, TRIM5α localizes at PML bodies when shuttling to the nucleus, as demonstrated by pharmacological treatment interfering with its nuclear export (43). The possibility of PML involvement in the inhibition of retroviruses by TRIM5α has been envisioned but not proven. The infectivity of the EIAV vector used here (which is restricted 5- to 10-fold by human TRIM5α (44)) was not significantly affected by knocking out PML (Fig. 2–4), suggesting that TRIM5α does not require PML. In order to increase sensitivity, we used an “N-tropic” strain of MLV, which is even more strongly restricted by human TRIM5α (39, 45) than EIAV, and of which restriction is counteracted by As_2_O_3_ in a cell context-specific fashion (46, 47). Thus, As_2_O_3_ greatly increases the infectivity of N-MLV but not B-MLV vectors in many human cell lines. The mechanism of action of As_2_O_3_ against TRIM5α has not been determined but it was thought to involve PML, since As_2_O_3_ is well-known as a specific inhibitor of PML (27, 48). Interestingly, As_2_O_3_ also enhances the infectivity of HIV-1 in human cells, although the magnitude of this effect is milder than what is found with N-MLV (46, 49). The molecular basis for the effect of As_2_O_3_ on HIV-1 and B-MLV vectors is unclear but is probably independent of TRIM5α and instead may be related to unidentified restriction factors (50, 51). We infected HeLa, TE671 and Jurkat cells with HIV-1_NL-GFP_, B-MLV_GFP_ and N-MLV_GFP_ in the presence of increasing As_2_O_3_ concentrations (Fig. 8A). In the absence of As_2_O_3_, N-MLV_GFP_ infectivity was barely detectable or undetectable in all three cell lines, reflecting the strong inhibition conferred by TRIM5α in human cells. At the same virus dose, B-MLV_GFP_ infected 3% to 5% of the cells.

**FIG 8.**
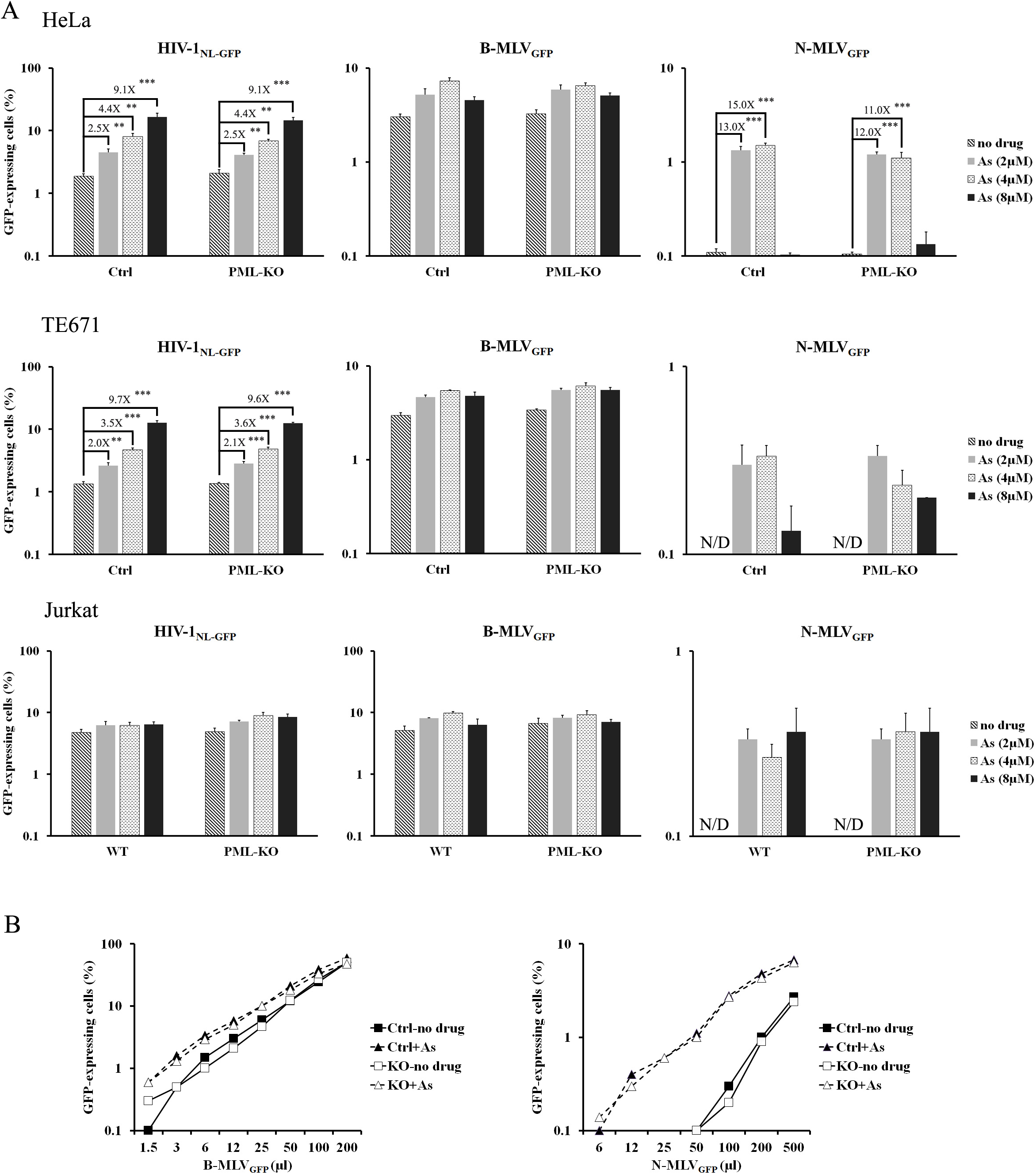
PML is irrelevant for the As_2_O_3_-induced stimulation of retroviral infectivity in human cells. (A) Effect of As_2_O_3_ on the permissiveness to retroviral vectors in the presence or absence of PML. Control and PML-KO human cell lines were treated with the indicated amounts of As_2_O_3_ for 15 min prior to infection with HIV-1, B-MLV and N-MLV vectors expressing GFP (B-MLV_GFP_ and N-MLV_GFP_ have identical titers in non-restrictive CRFK cat cells). The percentage of infected cells was assessed 2 d later by FACS. The values represent the means of three independent infections with standard deviations (N/D, not detected). Statistical significance was analyzed used a two-tailed Student's t-test (**P < 0.01, ***P < 0.001). (B) Virus dose-dependent infections. Ctrl and PML-KO HeLa cells were infected with increasing doses of B-MLV_GFP_ or N-MLV_GFP_ vectors in the presence or not of 4 µM As_2_O_3_. 2 d later, the percentage of infected cells was determined with FACS.

PML knockout had no effect on the infectivity of the two MLV vectors, implying that PML is not required for TRIM5α-mediated restriction of N-MLV. In presence of As_2_O_3_, N-MLV_GFP_ infectivity was greatly enhanced, although the stimulating effect was partly reversed at high As_2_O_3_ concentrations in HeLa and TE671 cells (Fig. 8A). As_2_O_3_ effectiveness at counteracting TRIM5α-mediated restrictions was found to decrease at high concentrations in previous studies as well (47, 49). In contrast to N-MLV_GFP_, B-MLV_GFP_ was only slightly enhanced by As_2_O_3_. As reported before, As_2_O_3_ modestly increased HIV-1_NL-GFP_ infection of HeLa and TE671 cells, although it had no effect on this vector in Jurkat cells (Fig. 8B). Knocking out PML had no detectable effect on the As_2_O_3_-mediated stimulation of N-MLV_GFP_ and HIV-1_NL-GFP_ in the three cell lines tested. We performed an additional infection of the HeLa cells with the N-MLV and B-MLV vectors, this time at a fixed As_2_O_3_ concentration and varying virus doses. Again, we observed that (i) PML had no effect on the infectivity of N-MLV_GFP_ and B-MLV_GFP_, (ii) As_2_O_3_-mediated stimulation of N-MLV_GFP_ was significantly stronger than that of B-MLV_GFP_, regardless of the virus dose, and (iii) knocking out PML had no impact on the effect of As_2_O_3_ on the MLV vectors. These data demonstrate that PML is not involved in the restriction of N-MLV by TRIM5α, nor is it involved in the mechanism by which As_2_O_3_ stimulates retroviral infections and counteracts TRIM5α

### PML is not required for TRIM5α-mediated restriction of HIV-1 in MEFs

MEFs provide a cellular environment in which PML restricts HIV-1, as seen by several laboratories (32–34). In addition, PML also inhibits HIV-1 transcription in MEFs, an effect that we did not observe in human cells (34). Thus, it would be conceivable for PML to have an impact on TRIM5α mediated restriction of HIV-1 in this specific cellular environment. To test this hypothesis, we used PML-KO MEFs (34, 52). WT and PML-KO MEFs were stably transduced with the HIV-1-restrictive Rhesus macaque TRIM5α or the non-restrictive human TRIM5α as a control. The cells were also transduced with the C35A RING domain mutant of each TRIM5α ortholog, which abolishes the RING domain-associated ubiquitin ligase activity (53). WB analyses showed that the transduced TRIM5α variants were expressed at comparable levels (Fig. 9A).

Colocalization of a fraction of TRIM5α with PML NBs was seen in the presence of the nuclear export inhibitor leptomycin B, consistent with published data obtained in human and canine cells (43), and exposure of the cells to HIV-1 did not modify this pattern (Fig. 9B). The cells were then challenged with HIV-1_NL-GFP_ or with the relatively restriction insensitive SIV_mac-GFP_ as a control (54), using virus doses at which PML has only mild effects on transduction by these lentiviral vectors in the absence of TRIM5α (34). HIV-1 was very strongly restricted by rhTRIM5α in both WT and PML-KO MEF cells (Fig. 9C). As expected, C35A rhTRIM5α and hTRIM5α (WT or C35A) had little to no effect on HIV-1_NL-GFP_, although we observed slightly higher levels of HIV-1 restriction by C35A rhTRIM5 in the presence of PML, perhaps suggesting that the presence of PML could partially compensate for the loss of a functional TRIM5α RING domain. SIV_mac-GFP_ was moderately restricted by rhTRIM5α, and PML knockout did not affect this inhibitory effect (in fact, restriction was slightly greater in the absence of PML) (Fig. 9C). In conclusion, PML is not required for rhTRIM5α to restrict HIV-1.

**FIG 9.**
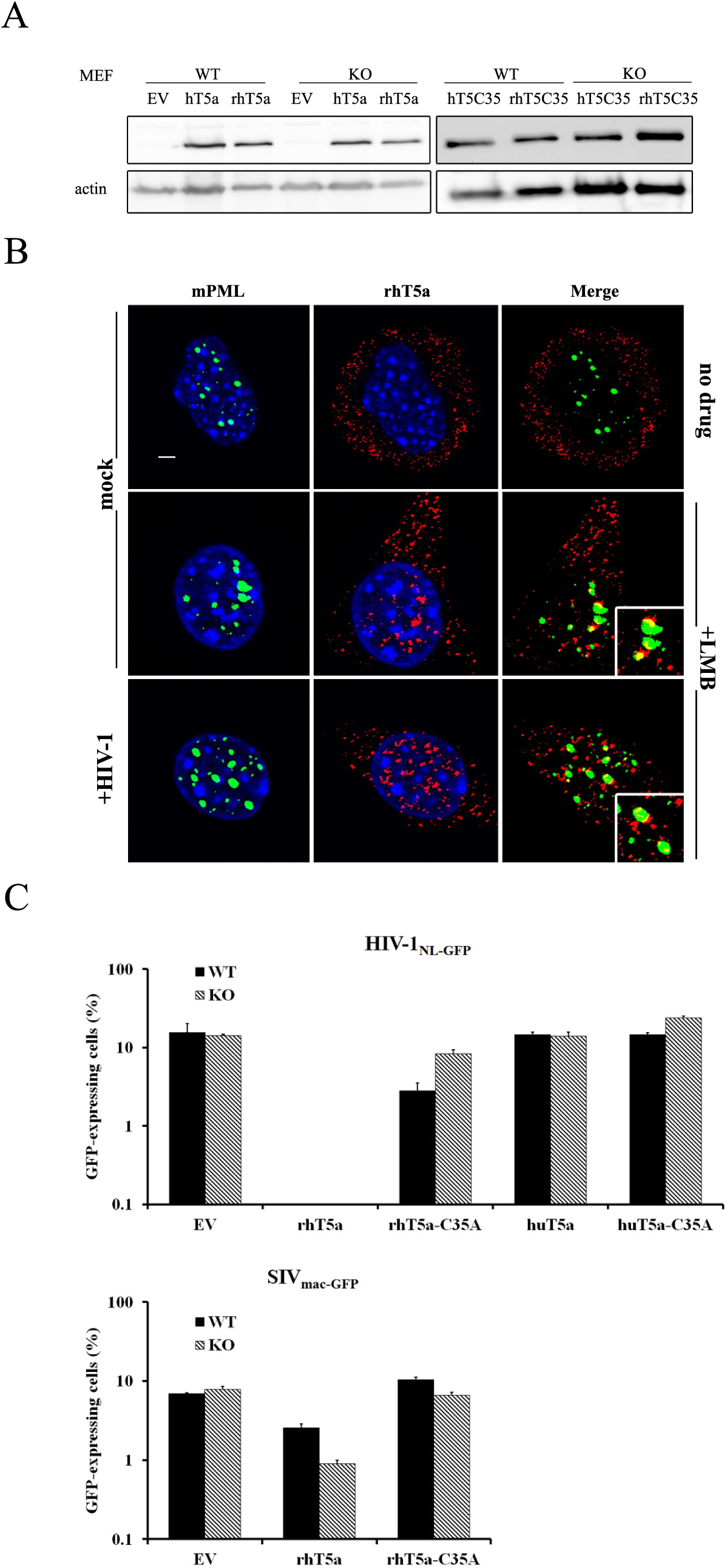
PML is not required for the TRIM5α-mediated restriction of HIV-1 in MEFs. (A) WB analysis of WT and mutant TRIM5α. MEFs were transduced with retroviral vectors expressing WT and C35A variants of FLAG-tagged rhTRIM5α and hTRIM5α. Following puromycin selection, cell lysates were prepared from the various cell populations transduced with the indicated vectors or transduced with the empty vector (EV) as a control. TRIM5α was detected using an antibody against FLAG, with actin used as a loading control. (B) Immunofluorescence staining of mPML and rhTRIM5α in WT MEFs stably transduced with FLAG-tagged rhTRIM5α. The cells were treated with either LMB (20 ng/ml) or PBS as a control, 3 h prior to infection with HIV-1_NL-GFP_ at a viral dose leading to approximately 10% infected cells. 6 h later, the cells were analyzed by immunofluorescence microscopy using anti-FLAG (red) and anti-mPML (green) antibodies. Nuclear DNA was stained using Hoechst 33342 (blue). Images are representative of multiple observations. Scale bar, 5 µm. (C) rhTRIM5α restricts HIV-1 in the presence or absence of PML. PML-KO or WT MEFs stably transduced with rhTRIM5α or huTRIM5α (WT or C35A mutant) were infected with HIV-1_NL-GFP_ or SIV_mac-GFP_, using virus amounts leading to infection of about 10% of the parental cells. 2 d later, the percentage of infected cells was measured by FACS. The values represent the means of three independent infections with standard deviations.

## Discussion

Whether PML has an impact or not on the infection of human cells by HIV-1 has been an open question for over 15 years. Trono and colleagues reported that PML is transiently exported in the cytoplasm following exposure to HIV-1 and co-localizes with the incoming virus in HeLa cells (55); however, this study did not include functional evidence for the involvement of PML in HIV-1 infection. Another team found no effect of HIV-1 infection on the distribution of PML bodies (56). As_2_O_3_, a known PML inhibitor, was found to enhance the infection of human cells with HIV-1 (46, 55) but it also stimulated the infection of MEFs with HIV-1 vectors whether PML was present or not (46). Interest for PML as a modulator of HIV-1 infection surfaced again in recent years, as it was proposed to act as an HIV-1 restriction factor in mouse and human cells (33). However, the data gathered so far by three different teams, including this study, suggest that the restriction activity in human cells, if it exists, is cell type-specific. Dutrieux and colleagues, using shRNAs, observed a modest inhibition of HIV-1 vector transduction conferred by PML in HeLa cells (<2-fold). They also observed a small delay in HIV-1 propagation in peripheral blood mononuclear cells, but the decrease in infectivity was not quantified (33). We previously observed that knocking down PML in T-lymphoid Sup-T1 cells increases HIV-1 infectivity by 2- to 4-fold (34). On the other hand, Kahle and colleagues saw no effect of knocking down PML on the infectivity of an HIV-1 vector in T lymphoid cell lines including CEM, HuT78, Jurkat and Molt4 (32). They showed, however, that PML reduces HIV-1 infectivity in human foreskin fibroblasts by 2- to 3-fold (32). Taken together, those previous papers showed that knocking down PML has either no effect or modest effects on HIV-1 infectivity in human cells. We were not able to efficiently knock out PML in Sup-T1 cells, preventing us from drawing comparisons with our previous knockdown results. However, our knockout experiments in Jurkat, THP-1, HeLa and TE671 are not consistent with PML being an HIV-1 restriction factor in human cells.

A recent study by Kim and Ahn (38) uncovered an additional function for PML in human skin fibroblasts: the stimulation of ISG expression through a direct association with their promoter. Accordingly, we previously showed that PML was important for the efficient inhibition of HIV-1 by IFN-I in MEF cells (34). Although HIV-1 is also readily inhibited by IFN-I in a variety of human cell types, as illustrated in our study, we find that this effect is not affected by knocking out PML. However, we cannot exclude the possibility that PML is involved in regulating IFN-I-dependent transcription in specific cellular contexts such as skin fibroblasts (38). It is also possible that PML stimulates the transcription of some ISGs but not others. In support of this idea is our observation that SIVmac inhibition by IFN-I in Jurkat cells was significantly greater in the presence of PML. SIVmac, but not HIV-1, is inhibited by an unidentified restriction factor in Jurkat cells and other T cells, provisionally called Lv4 (51). It is conceivable that the gene encoding Lv4 is specifically stimulated by IFN-I in a PML-dependent fashion in Jurkat cells. This characteristic could be exploited to identify this gene, similar to the strategy that led to the identification of Tetherin as a retroviral restriction factor (57).

In our previous study (34), we showed that PML inhibited HIV-1 transcription in MEFs but not in Sup-T1 cells and in an IFN-I-independent fashion. We analyzed GFP mean fluorescence intensity in all our experiments for this study, as a surrogate for HIV-1 gene expression levels. Consistent with our previous findings, we observed no effect of PML on the GFP fluorescence intensity following infection of THP-1, Jurkat, HeLa or TE671 cells with our various vectors (not shown). We conclude that PML does not repress HIV-1 transcription in human cells. This apparently contradicts a report by Giacca and colleagues that PML inhibits HIV-1 transcription by directly binding the viral promoter (31). However, the latter study was based on the use of “J-Lat” clones, which are Jurkat cells in which the HIV-1 provirus has become constitutively repressed through unknown mechanisms (58). We propose that PML may be involved in the rare silencing events leading to HIV-1 latency in Jurkat cells, and that PML is important for the maintenance of silencing; however, PML is not a ubiquitous silencer of HIV-1 transcription.

Finally, our study shows that the As_2_O_3_-mediated stimulation of early retroviral infection stages is completely independent of PML, and so is the inhibition of TRIM5α by this drug. Our experimental system was tailored to study the effect of As_2_O_3_ on restriction by TRIM5α and we cannot exclude that PML might be involved in other restriction activities known to be counteracted by As_2_O_3_ (50, 51). It is not entirely surprising that As_2_O_3_ inhibits TRIM5α in the absence of PML, considering that TRIM5α could target N-MLV in human cells, and HIV-1 in MEF cells, in the absence of PML. However, these results challenge conclusions from another paper that used radioactively or chemically labelled arsenate compounds to show that PML was the main target for this group of pharmacological agents (27). How, then, does As_2_O_3_ counteract TRIM5α and, to a lesser extent, stimulate HIV-1 and B-MLV vectors in human cells? Perhaps addressing this long-unanswered question will be helped by an observation that pre-dated the isolation of TRIM5α. Indeed, PK11195, a compound which, like As_2_O_3_, affects mitochondrial functions, also counteracts TRIM5α (49). Strikingly, these two drugs enhance autophagy (59, 60), an outcome possibly related to their effect on mitochondria. It is possible that As_2_O_3_-induced autophagy accelerates the lysosomal degradation of TRIM5α and other cytoplasmic restriction factors.

## Materials and methods

### Cell culture

Jurkat and THP-1 cells were maintained in RPMI 1640 medium (HyClone, Thermo Scientific, USA). Human embryonic kidney (HEK) 293T, HeLa, MEF and TE671 cells were maintained in Dulbecco's modified Eagle's medium (DMEM; HyClone). All culture media were supplemented with 10% fetal bovine serum (FBS) and penicillin/streptomycin (HyClone).

### Plasmids and preparation of retroviral vectors

The pMIP retroviral vector plasmids containing individual isoforms of hPML (pMIP-hPML-I to VI) and the isoform 2 of mPML (pMIP-mPML) have been described in details in a recent publication (34) and make use of materials generously provided by Roger D. Everett (61). Retroviral vectors were prepared by co-transfection of 293T cells with pMIP-m(h)PMLs together with pMD-G and pCL-Eco using polyethylenimine (PEI; Polyscience, Niles, IL) as detailed previously (34). Virus-containing supernatants were collected 2 d later, clarified by low-speed centrifugation and kept at −80 °C.

To produce GFP-expressing retroviral vectors, 293T cells were seeded in 10 cm culture dishes and transiently co-transfected with the following plasmids: pMD-G, pCNCG and pCIG3-B or pCIG3-N to produce B-MLV_GFP_ and N-MLV_GFP_, respectively; pMD-G and pHIV-1_NL-GFP_ to produce HIV-1_NL-GFP_; pMD-G and pSIV_mac239-GFP_ to produce SIV_mac-GFP_; or pONY3.1, pONY8.0 and pMD-G to produce EIAV_GFP_ (see (34, 62) and references therein).

### Design of gRNAs and transduction of lentiviral CRISPR-Cas9 vectors

The lentiviral expression vector plentiCRISPRv2 (pLCv2) was a gift from Feng Zhang (Addgene plasmid # 52961) and can be used to simultaneously express a gRNA, Cas9 nuclease, and puromycin resistance, either by transfection or lentiviral transduction (63). Two gRNAs (hPML1 and hPML2) targeting *hPML* (NG_029036) were designed using the Zhang lab online software available at crispr.mit.edu. The sequences targeted are 5'CAATCTGCCGGTACACCGAC (hPML1) and 5'CACCGGGAACTCCTCCTCCGAAGCG (hPML2). A gRNA targeting the

CAG hybrid promoter (target: 5'GTTCCGCGTTACATAACTTA) was used as a negative control (35). The oligodeoxynucleotides (ODNs) needed for the generation of pLCv2-based constructs were designed according to the Zhang lab protocol (63, 64) and are shown in Table S1.

The lentiviral vectors were prepared by co-transfection of 293T cells with 10 µg of the plentiCRISPRv2 construct together with 5 µg of pMD-G and 10 µg of pΔR8.9 (65). The viral supernatants were collected at 1.5 or 2 d post-transfection and used to transduce various cell lines. Stably transduced cells were selected by addition of 0.5 µg/ml puromycin (Thermo Fisher Scientific) to the medium at 2 d post-infection and for 5 d. Control untransduced cells were killed under these conditions.

### Surveyor nuclease and TIDE assays

To evaluate on-target modifications (indels) in *hPML*, a surveyor nuclease assay was performed. 293T cells were transfected with either plentiCRISPRv2-hPML1, -hPML2 or -CAG using PEI. 3 d later, the genomic DNA was extracted from the transfected cells using the QIAamp DNA mini kit (Qiagen, CA, USA). Two pairs of primers were designed to amplify 637 bp and 725 bp fragments on either side of Cas9 targets guided by gPML1 and gPML2 respectively (Fig. 1A). The sequences of these ODNs are included in Table S1. PCR amplicons were heat-denatured at 95 °C, and re-annealed by slow cooling to promote formation of dsDNA heteroduplexes. The heteroduplexes were then cleaved by surveyor nuclease S (Integrated DNA Technologies, Coralville, IA), according to the manufacturer's instructions. Digestion products were visualized by agarose gel electrophoresis. Amplicons containing the gPML1 target site were obtained from cells transduced with the lentiviral CRISPR vectors expressing gRNAs targeting hPML1 or CAG, using the WT/indel ODN pairs (see Table). These amplicons were Sanger sequenced using the WT/indel fwd ODN. A ∼175-nt long fragment of the sequencing data was then fed into the online TIDE assay that quantitates the % of indels by sequencing decomposition, in comparison with the unedited control (66).

**Table 1.**
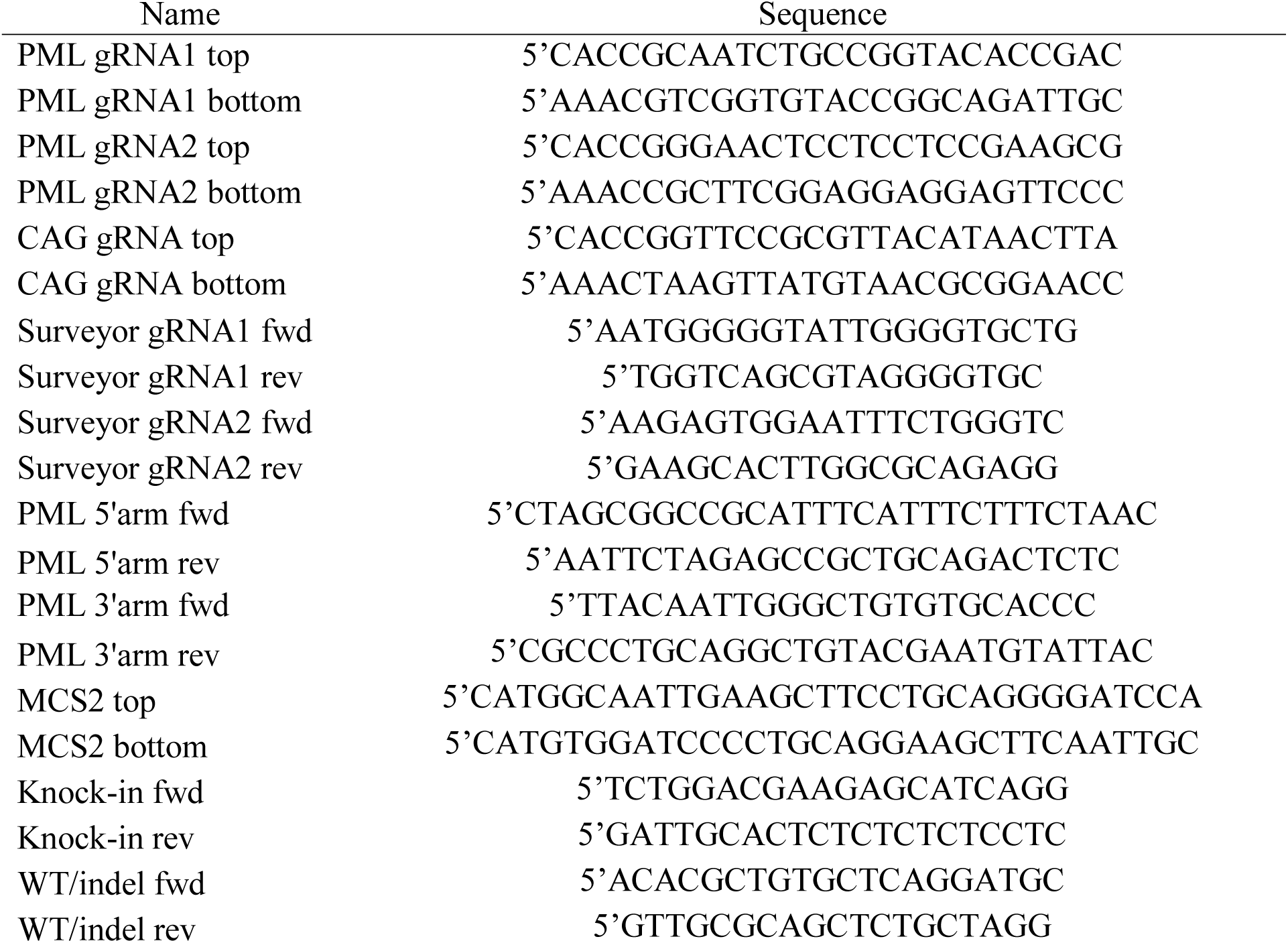
Sequence of ODNs used in this study.

### Construction of the homology directed repair (HDR) plasmid and generation of PML-KO Jurkat cells

We used pcDNA3.1+ as the backbone plasmid to prepare a HDR “donor” plasmid containing a neomycin selection gene (Neo^R^). First, the backbone plasmid was cut with BamHI and BglII, then self-ligated in order to remove the cytomegalovirus promoter from upstream of the multicloning site MCS1. Next, two ODNs were designed to introduce the second MCS (MCS2) (see Table S1); these ODNs were annealed, and the resulting duplex ligated into the PciI cut site of the plasmid, downstream of Neo^R^, yielding pNMs-Neo.HDR. To construct the PML HDR plasmid, homology arms corresponding to 800 bp-long regions immediately upstream and downstream of the hPML gRNA1-mediated Cas9 cut site in *hPML* were designed. The arms were amplified by PCR from genomic DNA extracted from 293T cells using the QIAamp DNA mini kit (Qiagen). The sequences of ODNs used in the PCR reactions are provided in Table S1. The 5’ arm was cloned into MCS1 of pNMs-Neo.HDR which had been cut with NotI and XbaI. The plasmid was then cut with Mfe I and Sbf I in order to clone the 3’ arm into MCS2, yielding pNMs-Neo.HDR-hPML.

Jurkat cells (300,000) were electroporated with 1.5 µg of pNMs-Neo.HDR-hPML together with 1.5 µg of pLCv2-hPML1 using an MP-100 microporator (Digital Bio Technology) according to the manufacturer's instructions. The parameters were 1300 V, 2 pulses, 20 ms. 48 h later, cells were placed in medium containing 1 mg/ml G418, and selection was carried out for 7 d.

#### Antibodies and WB analyses

Cells (1 × 10^6^) were lysed at 4°C in RIPA lysis buffer (1% NP40, 0.5% deoxycholate, 0.1% SDS, 150mM NaCl, 50 mM Tris-HCl pH 8.0). The lysates were subjected to SDS-polyacrylamide gel electrophoresis, followed by WB analysis using mouse anti-mPML mAb (36-1-104, Enzo life sciences, NY), rabbit polyclonal anti-hPML (A301-167A, Bethyl Laboratories, TX), rabbit polyclonal anti-FLAG (Cell Signaling, MA, USA), or mouse anti-β -actin antibody (Sigma, MI).

#### Viral challenges and flow cytometric analysis

Cells were seeded into 24-well plates at 3×10^4^ cells/well and infected the following day with GFP-expressing retroviral vectors. HeLa and TE671 cells were trypsinized at 2 d post-infection and fixed in 3% formaldehyde (Fisher Scientific, MA, USA). The percentage of GFP-positive cells was then determined by analyzing 1 × 10^4^ to 5 × 10^4^ cells on a FC500 MPL cytometer (Beckman Coulter, CA, USA) using the CXP Software (Beckman Coulter). All infection experiments were performed twice with identical results. One of two experiments is shown.

#### Pharmacological treatments

A 0.1 M stock solution of As_2_O_3_ (Sigma) was prepared in 1 N NaOH, as previously described (28), and diluted in the culture medium immediately before use. Cells were treated for 15 min prior to infection. 16 h post-infection, the supernatants were replaced with fresh medium devoid of drug. Recombinant human IFN- α was obtained from Shenandoah biotechnology (Warwick, PA). Recombinant human IFN- β and IFN-ω were obtained from PeproTech (Rocky Hill, NJ). IFN-I was added to cell cultures 16 h prior to infection and at a final concentration of 10 ng/ml.

#### Immunofluorescence microscopy

HeLa or MEF cells were seeded on glass coverslips placed in 3.5-cm wells. MEFs were treated with LMB (20 ng/ml) 3 h prior to infection then infected for 6 h with HIV-1_NL-GFP_. The cells were permeabilized and fixed for 10 min in Triton X-100/4% formaldehyde at room temperature (RT), followed by 4 washes with PBS. Cells were then treated with 10% goat serum (Sigma) for 30 min at RT followed by 4 h of incubation with antibodies against FLAG (Sigma, 1:150), hPML (Bethyl Laboratories, 1:150) or mPML (Enzo Life Sciences, 1:150) in 10% goat serum at RT. They were then washed 4 times with PBS and fluorescently stained with Alexa Fluor 488-conjugated goat anti-mouse or 594-conjugated goat anti-rabbit (Molecular Probes, Eugene, OR) diluted 1:100 in 10% goat serum for 1 h at RT. The cells were then washed 4 times with PBS before mounting in Vectashield (Vector Laboratories, Peterborough, UK). Hoechst 33342 (0.8 μg/ml; Molecular Probes) was added along with the penultimate PBS wash to reveal DNA. Images were acquired on an AxioObserver Microscope (Carl Zeiss Canada, Toronto, ON) equipped with the Apotome module.

## Acknowledgements

We thank Feng Zhang and Roger D. Everett for sharing reagents. This research received no specific grant from any funding agency in the public, commercial, or not-for-profit sectors.

